# A systematic review of delayed cerebral ischemia in experimental mouse models after subarachnoid hemorrhage

**DOI:** 10.1101/2023.12.28.573499

**Authors:** Cihat Karadag, Igor Fischer, Kimberley E. Wever, On Ying Chan, Rene Aquarius, Ronald H. M. A. Bartels, Jan F. Cornelius, Jasper H. van Lieshout, Hieronymus D. Boogaarts, Marcel A. Kamp

## Abstract

**Background:** The occurrence of delayed cerebral ischemia after aneurysmal subarachnoid hemorrhage is a main determinant for functional outcome. Despite two decades of animal studies investigating novel treatment strategies, the standard therapy for delayed cerebral ischemia has not changed. This translational gap raises the question to what extent experimental mouse models of subarachnoid hemorrhage can accurately mimic human delayed cerebral ischemia. The objective of this systematic review was to determine whether there are difference in the occurrence and timing of delayed cerebral ischemia between the various experimental mouse models of aneurysmal subarachnoid hemorrhage. Our aim was to identify which mouse model most closely mimics the human pathophysiology following aneurysmal subarachnoid hemorrhage.

**Methods:** This study was funded by ZonMw (project number 114024130). The review was preregistered at PROSPERO (protocol ID: CRD42020115578). A comprehensive search was performed in Medline via the PubMed interface and in EMBASE via the Ovid interface up to 2^nd^ November 2018The following exclusion criteria were used in both the title and abstract and full text phase: 1) not an original, full-length research paper, 2) no English language version available, 3) published before 1999, 4) not an animal study, 5) not a mouse study, 6) use of transgenic mice, 7) no subarachnoid hemorrhage induction, 8) no delayed cerebral ischemia measured, 9) reported an observation time less than six hours, 10) cross-over study, or any study design without a control group. Data was extracted by one reviewer. The SYRCLE risk of bias tool was used in duplo by two independent reviewers to assess risks of bias in the included studies, with discrepancies being resolved through discussion. A narrative synthesis of the evidence was performed.

**Results:** The literature search retrieved a total of 1461 papers, of which 71 publications met the inclusion criteria. Most studies were assessed at an unclear risk for most types of bias. Mice models were highly standardized: the C57Bl/6 strain was used in 53 studies (74.6%), only male animals were used in 55 studies (77.5%). To model a subarachnoid hemorrhage, perforation of the anterior cerebral artery / internal carotid artery with a suture was performed in 43 studies (60.6%), while direct injection of blood was performed in 24 studies (33.8%). The presence of delayed cerebral ischemia was established through neurological outcomes (44 studies, 62.0%), ex-vivo histology (39 studies, 54.9%) or in-vivo imaging (10 studies, 14.1%) assessment. Incidence of delayed cerebral ischemia was similarly high for all outcome measures (between 85-87%) and the timing of delayed cerebral ischemia occurrence was similar, despite differences in subarachnoid hemorrhage induction method.

**Conclusion:** Our results show that established perforation and injection-models for experimental subarachnoid hemorrhage are highly standardized. Yet, there is a high inconsistency in definitions of delayed cerebral ischemia in experimental mouse models. Although perforation model results in a higher rate of delayed cerebral ischemia, there is no effect on the time of occurrence after ictus. It is unclear to what extent the delayed cerebral ischemia demonstrated in these animal models is comparable to the clinical situation.

## Introduction

Delayed cerebral ischemia is a main determinant for functional outcome after aneurysmal subarachnoid hemorrhage [79,42]. The prevalence is 0.29% and remains constant over the last decades [65]. It has been well defined in humans and has been studied extensively in order to treat related symptoms, determine the prognosis of patients and improve functional outcome after aneurysmal subarachnoid hemorrhage [79,83].

Animal models are useful to study drug safety, enable researchers to analyze complex pathophysiological and molecular mechanisms and provide ample options for genetic engineering. Mice are the most common laboratory animals, because of the great knowledge about their physiology and the availability of technical resources and transgenic animals [23]. They were the latest species that was introduced into experimental subarachnoid hemorrhage research in 1999 [44]. The number of studies in mice has steadily increased during the last two decades and has - together with rat and rabbit models - become one of the most commonly used species in preclinical SAH research [45]. Various pharmaceutics for the treatment of delayed cerebral ischemia showed promising results in *in vivo* mouse studies [28]. However, despite the high number of studies, different therapeutic approaches and numerous analyzed pharmaceutics, standard therapy for delayed cerebral ischemia has not changed.

This translational gap raises the question to what extent experimental mouse models can mimic human aneurysmal subarachnoid hemorrhage. Mice differ from humans in cerebral anatomy, anatomy of the cerebral vessels and hemodynamics, pathophysiology and metabolism [29]. Moreover, the genetic background of mice influences the occurrence and severity of delayed cerebral ischemia and the selected SAH model affects delayed cerebral ischemia-associated mortality after experimental subarachnoid hemorrhage in mice [27,10,11]. Still, potential influence of the model used on the occurrence and timing of delayed cerebral ischemia has not yet been investigated.

The aim of this systematic and meta-analysis is therefore to evaluate if the occurrence and timing of delayed cerebral ischemia differs between experimental mouse models of aneurysmal subarachnoid hemorrhage. Our aim was to identify which model most closely mimics the human pathophysiology following aneurysmal subarachnoid hemorrhage.

## Methods

### Reporting and protocol

This report follows the PRISMA guidelines for systematic reviews. The review methodology was recorded *a priori* in accordance with SYRCLE’s systematic review protocol for animal intervention studies [12] and registered in the international prospective register of systematic reviews PROSPERO (protocol ID: CRD42020115578). No amendments to the review protocol were made during the study.

### Search strategy

A medical information specialist (O.C.) performed a comprehensive search in PubMed and Embase databases. The aim was to identify all studies ranging from the first description of a mouse SAH model in 1999 (first description of a SAH mouse model) up to 2^nd^ November 2018, for studies reporting on DCI-related pathophysiological changes in comparison to a control group. Detailed information on the search strategy is provided in Supplement 1. Moreover, we screened reference lists of all already included studies to identify additional studies eligible for inclusion.

### Study selection

After removal of duplicates using Endnote (X9.2, Clarivate Analytics, United States), the search results were imported into the online screening platform Rayyan (Rayyan.ai, Rayyan Systems Inc., United States). In the first phase, references were screened for eligibility by two reviewers (JvL and RA) based on title and abstract. Eligible papers were subsequently screened for final inclusion (by JvL and CK) based on the full text. In case of discrepancies, another independent reviewer was contacted in order to reach agreement (KW). In both phases, studies were excluded if at least one of the following exclusion criteria was applicable:

1. not a full research article with original data,
2. no English language version available,
3. published before 1999,
4. not an animal study,
5. not a mouse study,
6. use of transgenic mice,
7. not an aneurysmal subarachnoid hemorrhage model,
8. no delayed cerebral ischemia measured
9. reported an observation time less than six hours,
10. cross-over study, or any study design without a control group,

### Data extraction

Data extraction was performed by one author (CK). The following data was extracted from each study: bibliographic details (author, journal and year of publication), study details (number, type and size of the experimental and control groups), animal model details (strain, age, sex and weight), model induction details (SAH induction, SAH location, type of blood used, amount of blood injected, type of suture used and monitoring duration), detection of DCI (occurrence (Y/N), way of DCI detection (histology, imaging, functional)), measured time of onset of DCI.

### Outcome measures and data synthesis

Similarities and differences in animal models will be displayed as well as the occurrence of DCI and the way DCI was detected. No meta-analysis was planned.

### Assessment of methodological quality and risks of bias

The internal validity of the included studies was assessed according to SYRCLE’s risk of bias (RoB) tool for animal studies [24]. Two independent authors (CK, JvL) reviewed the included publications using this tool. Discrepancies were resolved through discussion.

## Results

### Included studies

The literature search revealed a total of 1461 records, 541 in PubMed, 913 in Embase. An additional seven studies were found by screening of the references. After removal of duplicates, we were able to identify 1008 studies. Screening the abstracts against the exclusion criteria yielded 138 studies eligible for full-text evaluation. Another 67 studies were excluded during full-text screening based upon our exclusion criteria. In total, we identified 71 studies that met our inclusion criteria (Figure 1).

**Figure 1:**
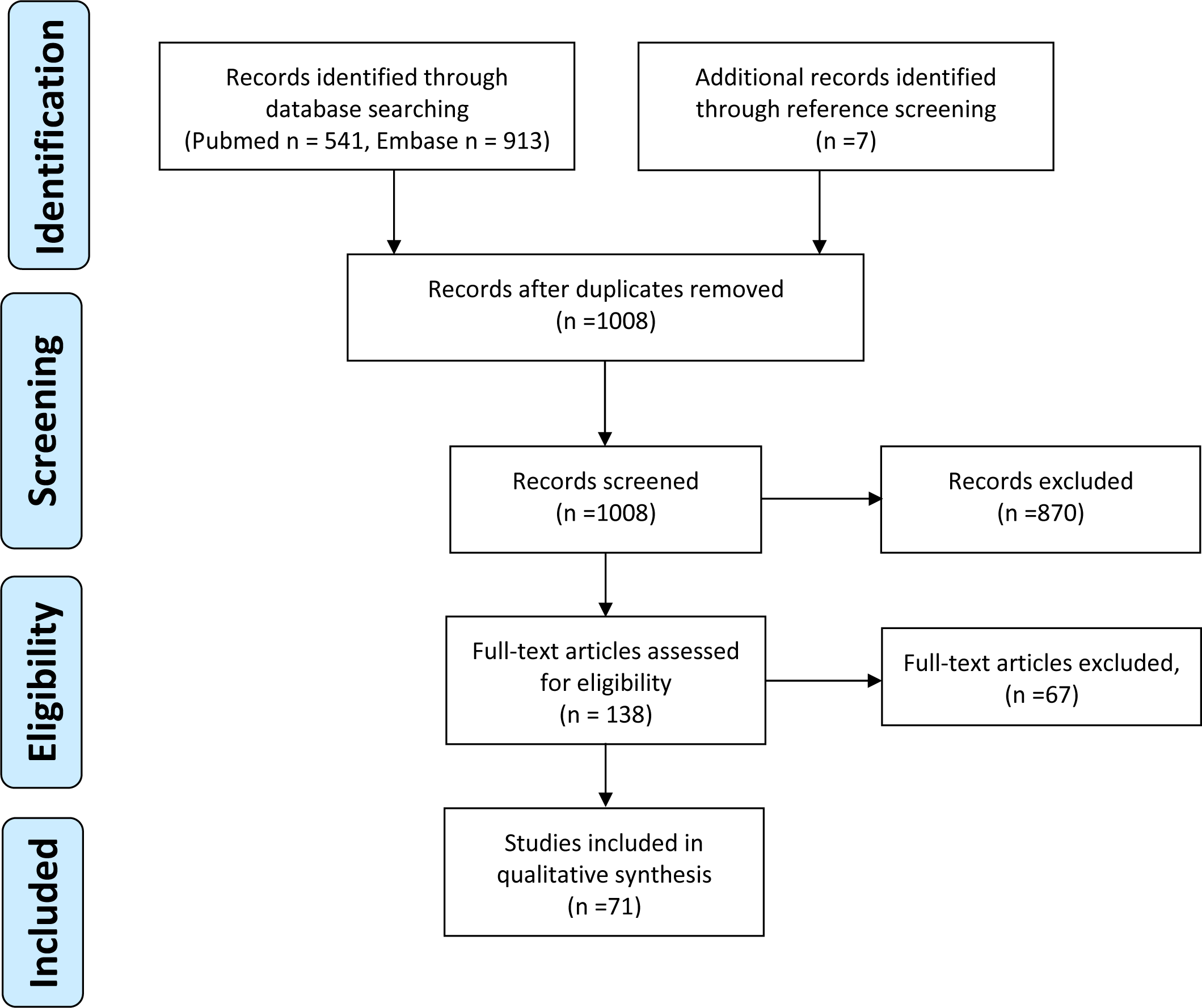
Prisma Flow Diagram

### Study characteristics

A total of 71 studies were included for data extraction and risk of bias analysis. See Table 1 and Supplement 2 for a full overview of the study characteristics.

| ***Study*** | ***Strain*** | ***Sex*** | ***Weight (g)*** | ***Age (weeks)*** | ***SAH induction method*** | ***SAH location*** | ***Type of blood*** | ***Injection Volume (μl)*** | ***Filament size*** |
| --- | --- | --- | --- | --- | --- | --- | --- | --- | --- |
| Altay2009 [4] | C57Bl/6 | Male | 28-32 | 10-12 | Other | CM | Venous | N/A | N/A |
| Altay2012 [3] | CD-1 | Male | 30-38 | 8 | Perforation | ACA/ICA | Arterial | N/A | 4-0 |
| Altay2012 [1] | CD-1 | Male | 30-38 | N/R | Perforation | ACA/ICA | Arterial | N/A | 4-0 |
| Altay2014 [2] | CD-1 | Male | 30-38 | 8 | Perforation | ACA/ICA | Arterial | N/A | 4-0 |
| Ayer2011 [5] | CD-1 | Male | 20-25 | N/R | Perforation | ACA/ICA | Arterial | N/A | 5-0 |
| Boettinger2017 [6] | C57Bl/6 | Both | 22-28 | 10-12 | Injection | PC | Arterial | 100-120 | N/A |
| Bühler2014 [7] | C57Bl/6 | Male | 22-25 | N/R | Perforation | ACA/ICA | Arterial | N/A | 5-0 |
| Chaichana2007 [8] | C57Bl/6 | Both | 22-27 | N/R | Injection | CM | Arterial | 60 | N/A |
| D'Abbodanza2015 [11] | Mixed | Male | 20-25 | N/R | Injection | PC | Arterial | 80 | N/A |
| D'Abbodanza2016 [10] | Mixed | Male | 20-25 | N/R | Injection | CM | Arterial | 60 | N/A |
| Dong2016 [13] | C57Bl/6 | Male | 26-30 | 10-12 | Perforation | ACA/ICA | Arterial | N/A | 5-0 |
| Du2016 [14] | C57Bl/6 | Male | 23-26 | N/R | Perforation | ACA/ICA | Arterial | N/A | 5-0 |
| El Amki2018 [15] | C57Bl/6 | N/R | 25-30 | 12-14 | Injection | CM | Arterial | 90 | N/A |
| Feiler2010 [16] | C57Bl/6 | Male | 23-25 | N/R | Perforation | ACA/ICA | Arterial | N/A | 5-0 |
| Friedrich2012 [17] | C57Bl/6 | Male | 23-25 | N/R | Perforation | ACA/ICA | Arterial | N/A | 5-0 |
| Fujimoto2016 [18] | C57Bl/6 | Male | 20-25 | N/R | Perforation | ACA/ICA | Arterial | N/A | 5-0 |
| Fujimoto2017 [30] | C57Bl/6 | Female | 20-25 | N/R | Perforation | ACA/ICA | Arterial | N/A | 5-0 |
| Gao2006 [19] | C57Bl/6 | Male | N/R | 12-14 | Perforation | ACA/ICA | Arterial | N/A | 5-0 |
| Garzon Muvdi2013 [20] | C57Bl/6 | N/R | 22-30 | N/R | Injection | CM | Arterial | 60 | N/A |
| Han2012 [21] | C57Bl/6 | Male | N/R | 12-16 | Perforation | ACA/ICA | Arterial | N/A | 5-0 |
| Hanafy2013 [22] | C57Bl/6 | Male | N/R | 10-12 | Injection | PC | Arterial | 60 | N/A |
| Kamii1999 [25] | N/R | N/R | N/R | N/R | Perforation | ACA/ICA | Arterial | N/A | 5-0 |
| Kamp2014 [26] | C57Bl/6 | Male | N/R | 15-20 | Injection | CM | Venous | 50 | N/A |
| Kawakita2017 [30] | C57Bl/6 | Male | 25-30 | N/R | Perforation | ACA/ICA | Arterial | N/A | 4-0 |
| LeBlanc2016 [31] | C57Bl/6 | Male | 24-27 | N/R | Injection | PC | Arterial | 60 | N/A |
| Li2014 [35] | ICR | Male | N/R | N/R | Injection | PC | Arterial | 100 | N/A |
| Li2015 [34] | ICR | Female | 30-35 | N/R | Injection | PC | Arterial | 100 | N/A |
| Li2016 [33] | C57Bl/6 | Male | 25-30 | N/R | Perforation | ACA/ICA | Arterial | N/A | 6-0 |
| Li2020 [36] | C57Bl/6 | Male | 20-25 | 8-12 | Perforation | ACA/ICA | Arterial | N/A | 5-0 |
| Lin2002 [37] | C57Bl/6 | Male | 27-32 | N/R | Injection | CM | Arterial | 60 | N/A |
| Lin2005 [38] | C57Bl/6 | Male | 30-35 | N/R | Injection | CM | Arterial | 400 | N/A |
| Liu2007 [40] | C57Bl/6 | N/R | N/R | 12-20 | Perforation | ACA/ICA | Arterial | N/A | 5-0 |
| Liu2016 [39] | C57Bl/6 | Female | 25-30 | 10-12 | Perforation | ACA/ICA | Arterial | N/A | 4-0 |
| Luo2016 [41] | C57Bl/6 | N/R | 20-25 | 8-12 | Injection | CM | Arterial | 60 | N/A |
| McConnell2016 [46] | FVB/Nj | Both | N/R | 8-12 | Injection | CM | Arterial | 60-80 | N/A |
| McGirt2002a [48] | C57Bl/6 | Male | N/R | 10-12 | Perforation | ACA/ICA | Arterial | N/A | 5-0 |
| McGirt2002b [47] | C57Bl/6 | Male | N/R | 10-12 | Perforation | ACA/ICA | Arterial | N/A | 5-0 |
| Milner2014 [49] | C57Bl/6 | Male | 24-30 | 12-14 | Perforation | ACA/ICA | Arterial | N/A | 5-0 |
| Milner2014 [50] | C57Bl/6 | Male | 24-30 | 12-14 | Perforation | ACA/ICA | Arterial | N/A | 5-0 |
| Mo2013 [51] | Other | Male | 18-20 | N/R | Injection | CM | OxyHb | 50 | N/A |
| Muroi2014 [53] | C57Bl/6 | Male | 21-26 | 10-12 | Perforation | ACA/ICA | Arterial | N/A | 5-0 |
| Muroi2014 [52] | C57Bl/6 | Male | 22-26 | N/R | Perforation | ACA/ICA | Arterial | N/A | 5-0 |
| Mutoh2017 [54] | C57Bl/6 | Male | 21-26 | 7 | Perforation | ACA/ICA | Arterial | N/A | 5-0 |
| Nakano2018 [55] | C57Bl/6 | Male | 25-30 | 10-12 | Perforation | ACA/ICA | Arterial | N/A | 4-0 |
| Neulen2017 [57] | C57Bl/6 | Both | N/R | 24-27 | Perforation | ACA/ICA | Arteria | N/A | 5-0 |
| Neulen2018 [56] | C57Bl/6 | Male | N/R | 10-12 | Perforation | ACA/ICA | Arterial | N/A | 5-0 |
| Neulen2019 [58] | C57Bl/6 | Male | N/R | 11-14 | Perforation | ACA/ICA | Arterial | N/A | 5-0 |
| Oka2017 [59] | C57Bl/6 | Male | 25-30 | 12-16 | Injection | PC | Arterial | 40 | N/A |
| Parra2002 [60] | C57Bl/6 | Both | N/R | 8-10 | Perforation | ACA/ICA | Arterial | N/A | 5-0 |
| Pradilla2011 [62] | C57Bl/6 | Both | 22-30 | N/R | Injection | CM | Arterial | 60 | N/A |
| Provencio2011 [63] | C57Bl/6 | Male | 20-32 | 10-12 | Other | CM | Venous | N/A | N/A |
| Provencio2016 [64] | C57Bl/6 | Male | N/R | 8-12 | Other | CM | Venous | N/A | N/A |
| Sabri2009 [67] | CD-1 | Male | 33-40 | N/R | Injection | PC | Arterial | 100 | N/A |
| Sabri2012 [66] | CD-1 | Male | 20-25 | N/R | Injection | PC | Arterial | 100 | N/A |
| Sheng2011 [69] | C57Bl/6 | Male | 20-25 | N/R | Perforation | ACA/ICA | Arterial | N/A | 5-0 |
| Sheng2011 [68] | C57Bl/6 | Male | 20-25 | N/R | Perforation | ACA/ICA | Arterial | N/A | 5-0 |
| Siler2014 [71] | C57Bl/6 | Male | N/R | 8-12 | Perforation | ACA/ICA | Arterial | N/A | 5-0 |
| Siler2015 [70] | C57Bl/6 | Male | N/R | 8-12 | Perforation | ACA/ICA | Arterial | N/A | 5-0 |
| Siler2015 [72] | C57Bl/6 | Male | N/R | 8-12 | Perforation | ACA/ICA | Arterial | N/A | 5-0 |
| Smithason2013 [73] | C57Bl/6 | Male | 20-32 | 10-12 | Other | CM | Venous | N/A | N/A |
| Suzuki2009 [74] | CD-1 | N/R | 35-40 | N/R | Perforation | ACA/ICA | Arterial | N/A | 4-0 |
| Tait2010 [75] | CD-1 | Male | 30-35 | 6-8 | Injection | PC | Arterial | 30 | N/A |
| Terpolilli2016 [76] | C57Bl/6 | Male | 24-26 | 6-8 | Perforation | ACA/ICA | Arterial | N/A | 5-0 |
| Tso2015 [77] | C57Bl/6 | Male | 20-25 | 8-10 | Injection | PC | Arterial | 80 | N/A |
| Turan2018 [78] | C57Bl/6 | Male | N/R | 6 | Injection | CM | Arterial | 60 | N/A |
| Vellimana2011 [81] | C57Bl/6 | Male | N/R | 12-16 | Perforation | ACA/ICA | Arterial | N/A | 5-0 |
| Vellimana2017 [82] | C57Bl/6 | Male | N/R | 12-14 | Perforation | ACA/ICA | Arterial | N/A | 5-0 |
| Wakade2009 [85] | CD-1 | Male | N/R | 8-10 | Perforation | ACA/ICA | Arterial | N/A | 5-0 |
| Xu 2015 [86] | ICR | Male | 28-32 | N/R | Perforation | ACA/ICA | Arterial | N/A | 5-0 |
| Yagi2015 [87] | C57Bl/6 | Male | N/R | N/R | Injection | PC | Arterial | 80 | N/A |
| Yeung2013 [88] | N/R | N/R | N/R | N/R | Perforation | ACA/ICA | Arterial | N/A | 5-0 |

N/A: Not applicable; N/R: Not reported; ACA: Anterior cerebral artery; ICA: Internal carotid artery; CM: Cisterna Magna; PC: prechiasmatic

Strains most used were C57Bl/6 (n=53, 74.6%) and CD-1 (n=9, 12.7%). Two studies (2.8%) did not report on the strain used for the described experiment. Male animals were used in 55 studies (77.5%), female animals were used in 3 studies (4.2%), both sexes were used in 6 studies (8.5%) and sex was not reported in 7 studies (9.9%). Weight of the mice used was <20g in 1 study (1.4%), 20-30g in 38 studies (53.5%) and >30 g in 8 studies (11.3%). Weight was not reported in 24 studies (33.8%). Age of mice used was <10 weeks in 19 studies (26.8%), 10-20 weeks in 20 studies (28.2%) and >20 weeks in 1 study (1.4%). Age was not reported in 31 studies (43.7%). Both weight and age were not reported in 4 studies (5.6%). Strain, sex, weight and age were not reported in 2 studies (2.8%).

Perforation of an intracranial vessel with a suture was performed in 43 studies (60.6%) In all perforation models, the anterior cerebral artery / internal carotid artery was the origin of the subarachnoid hemorrhage. The suture used to accomplish the SAH was 4-0 in 7 studies (16.3%), 5-0 in 35 studies (81.4%) and 6-0 in 1 study (2.3%). Direct injection of blood was performed in 24 studies (33.8%). Arterial blood was injected in most studies (n=22, 91.7%) and the injected volume differed considerably among studies: ≤50 µL was injected in 2 studies (8.3%), 50-100 µL was injected in 21 studies (87.5%), ≥100µL was injected in 1 studies (4.2%), namely 400 µL. Other methods to induce a SAH were chosen in 4 studies (5.6%).

### Quality assessment

By scoring the reporting or key quality indicaors, we demonstrated that randomization, blinding and conflicts of interest were mentioned in 49.3%, 76.1% and 63.4% of the included studies, respectively. A sample size calculation was mentioned in 12.7% of the studies (Figure 2 and Supplement 3).

**Figure 2:**
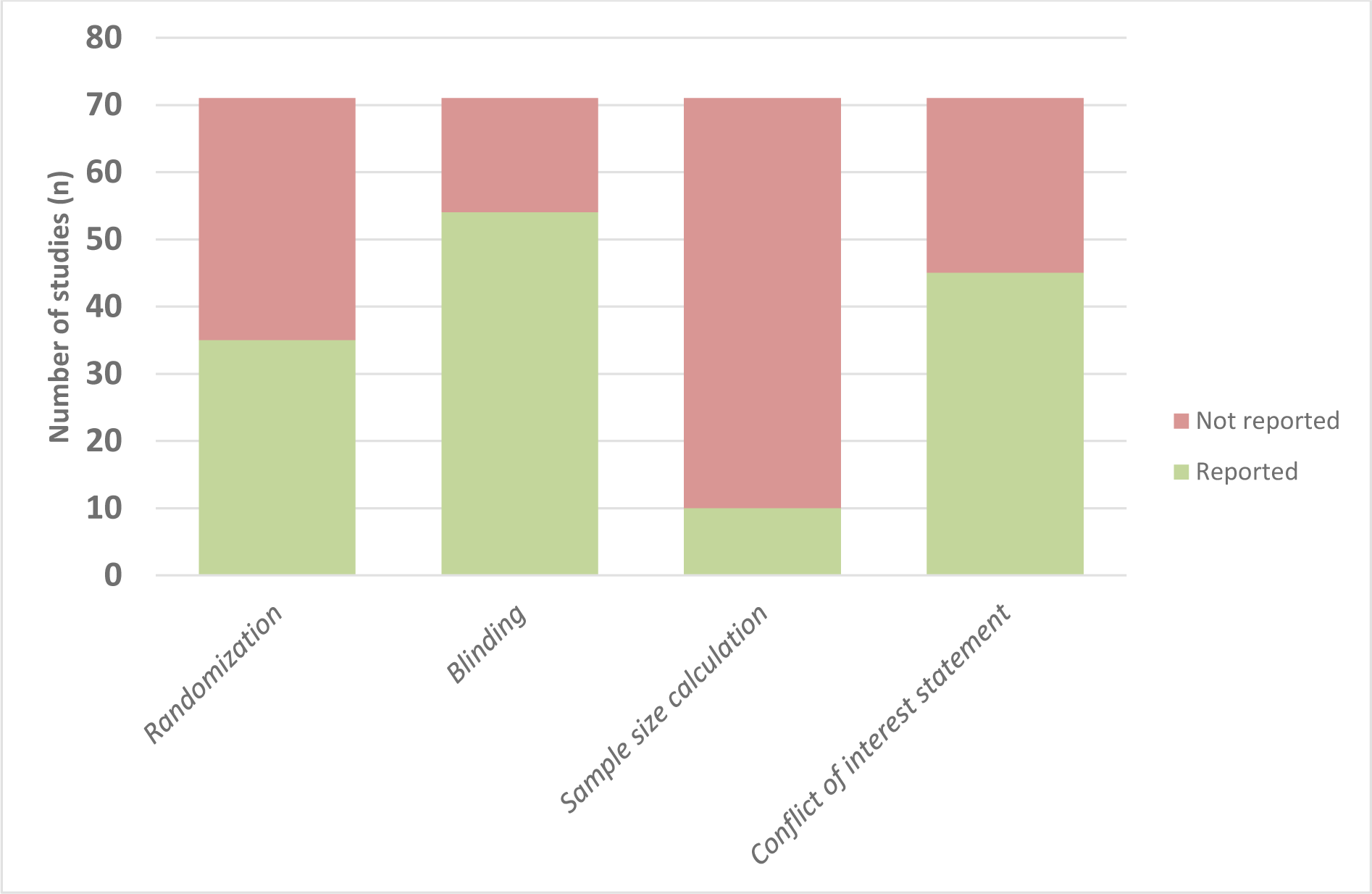
Frequency of reporting of 4 key study quality indicators.

The risk of bias assessment using SYRCLE’s risk of bias tool demonstrates that most studies were at an unclear risk for most biases (Figure 3 and Supplement 3). The only items for which >50% of the studies scored low risk of bias were selection bias as a result of baseline differences in characteristics between the groups (74.6%) and detection bias as a result of non-blinded outcome assessment (76.1%).

**Figure 3:**
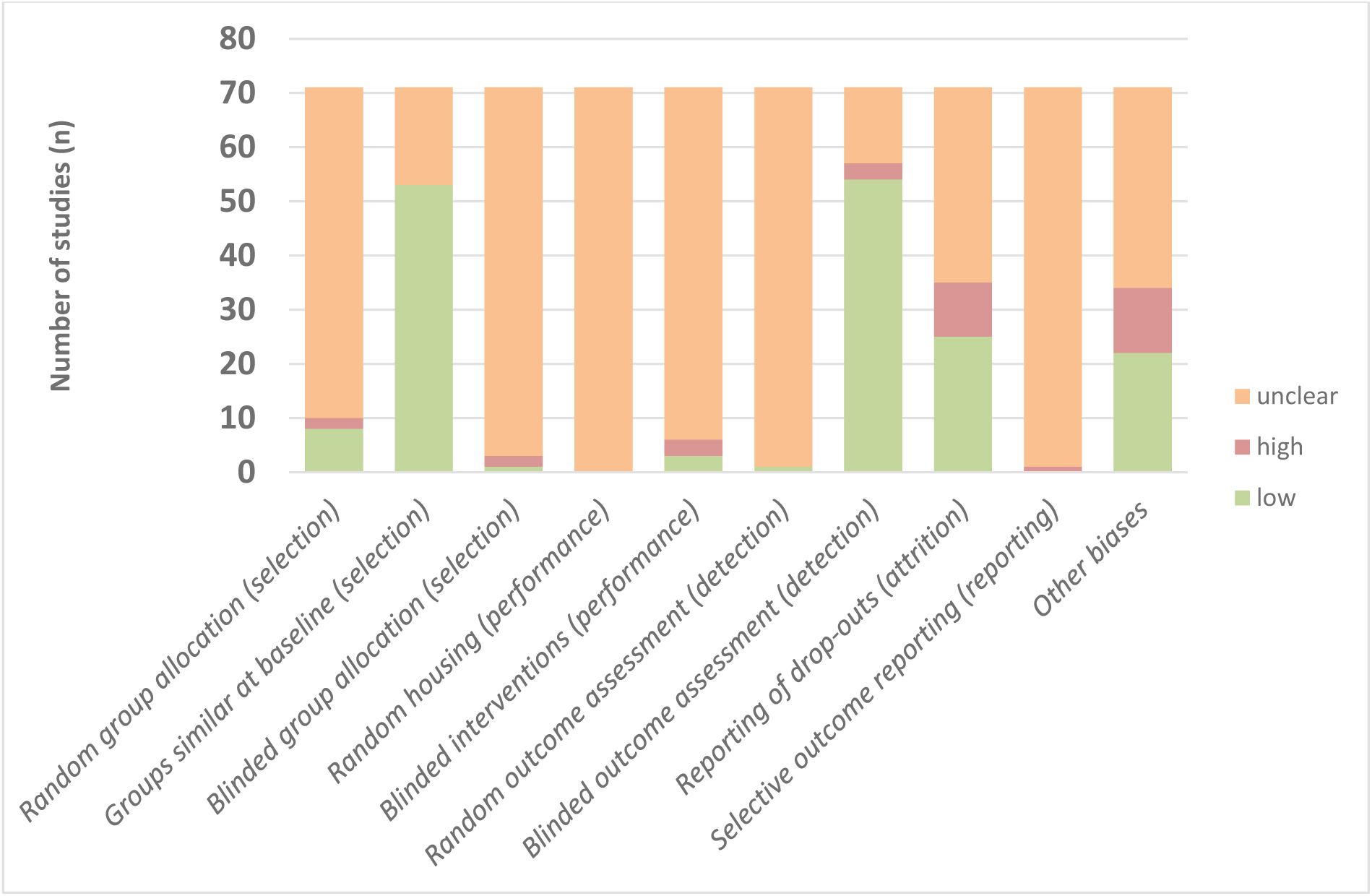
Visual representation of the SYRCLE risk of bias assessment.

### Data synthesis

Occurrence of delayed cerebral ischemia was most often confirmed by neurological outcome scores (44 studies, 62.0%), followed by ex-vivo (histological) examination (39 studies, 54.9%) and in-vivo imaging (10 studies, 14.1%).

It was possible to calculate the average percentage of animals affected by delayed cerebral ischemia after subarachnoid hemorrhage (at any time point) as determined through neurological (n=42 studies), histological (n=36 studies) or in-vivo image (n=8 studies) analysis. All relevant results are displayed in Table 2.

**Table 2:** Presence of delayed cerebral ischemia in mouse models of subarachnoid hemorrhage as described in studies.

|  | Presence of DCI in mouse models of SAH – irrespective of timing of assessment |  |  |  |  |
| --- | --- | --- | --- | --- | --- |
|  | Number of studies (n) | Average percentage of animals with DCI (%) | Standard deviation of animals with DCI | Minimum (%) | Maximum (%) |
| **Neurology** | 42 | 87.4 | 25.7 | 0 | 100 |
| **Histology** | 36 | 85.3 | 23.3 | 35 | 100 |
| **Imaging** | 8 | 86.4 | 27.0 | 27.3 | 100 |

DCI: delayed cerebral ischemia. SAH: subarachnoid hemorrhage

Timing of the occurrence of delayed cerebral ischemia was very heterogeneous among studies. Most studies assessed the occurrence of delayed cerebral ischemia at 24, 38 or 72 hours after ictus, but studies have measured delayed cerebral ischemia up to 336 hours after ictus (Table 3). The two most used induction methods of subarachnoid hemorrhage, arterial perforation and blood injection, were represented over the full width of measurement times after ictus (Table 3).

**Table 3:** Number of studies reporting the presence of delayed cerebral ischemia in mouse models with various induction methods of subarachnoid hemorrhage.

|  | Number of studies reporting the presence of DCI in various induction methods of SAH (n) |  |  |  |  |  |  |  |  |  |  |  |  |  |  |  |
| --- | --- | --- | --- | --- | --- | --- | --- | --- | --- | --- | --- | --- | --- | --- | --- | --- |
| *Time of Measurement (hours after ictus)* | 6 | 12 | 21 | 23 | 24 | 36 | 48 | 72 | 96 | 120 | 144 | 168 | 192 | 216 | 240 | 336 |
| **Neurology** | P:0<br>I:1<br>O:0 | P:0<br>I:1<br>O:0 | P:0<br>I:1<br>O:0 | P:1<br>I:0<br>O:0 | P:23<br>I:6<br>O:0 |  | P:13<br>I:2<br>O:0 | P:14<br>I:2<br>O:1 | P:2<br>I:1<br>O:0 | P:1<br>I:1<br>O:1 | P:1<br>I:1<br>O:1 | P:4<br>I:2<br>O:1 | P:0<br>I:0<br>O:1 | P:0<br>I:1<br>O:2 | P:0<br>I:1<br>O:1 | P:2<br>I:0<br>O:0 |
| **Histology** | P:0<br>I:1<br>O:0 | P:1<br>I:1<br>O:0 |  |  | P:5<br>I:6<br>O:2 | P:0<br>I:1<br>O:0 | P:4<br>I:3<br>O:0 | P:16<br>I:2<br>O:0 |  | P:0<br>I:1<br>O:0 | P:0<br>I:0<br>O:3 | P:0<br>I:4<br>O:0 |  |  | P:0<br>I:2<br>O:0 |  |
| **Imaging** | P:2<br>I:1<br>O:0 | P:1<br>I:2<br>O:0 |  |  | P:5<br>I:0<br>O:0 |  | P:1<br>I:2<br>O:0 | P:3<br>I:0<br>O:0 | P:1<br>I:0<br>O:0 | P:1<br>I:0<br>O:0 | P:1<br>I:0<br>O:0 |  |  |  |  |  |
| **Total** | P:2<br>I:3<br>O:0 | P:2<br>I:4<br>O:0 | P:0<br>I:1<br>O:0 | P:1<br>I:0<br>O:0 | P:33<br>I:12<br>O:2 | P:0<br>I:1<br>O:0 | P:18<br>I:7<br>O:0 | P:33<br>I:4<br>O:1 | P:3<br>I:1<br>O:0 | P:2<br>I:2<br>O:1 | P:2<br>I:1<br>O:4 | P:4<br>I:6<br>O:1 | P:3<br>I:1<br>O:1 | P:0<br>I:1<br>O:2 | P:0<br>I:3<br>O:1 | P:2<br>I:1<br>O:0 |

DCI: delayed cerebral ischemia. SAH: subarachnoid hemorrhage. P: Perforation model, I: Injection model, O: Other model.

## Discussion

In this systematic review we have created an overview of the literature to demonstrate the variation in the induction methods in mouse models of subarachnoid hemorrhage and the timing in the presence of delayed cerebral ischemia.

The first observation is that most experiments observed delayed cerebral ischemia at 24, 48 and 72 hours after subarachnoid hemorrhage induction. In humans, all of these time points would fall under early brain injury, the precursor of delayed cerebral ischemia [9]. However, it has never been defined where early brain injury stops and delayed cerebral ischemia begins in mouse models of experimental subarachnoid hemorrhage. Recently, scientists have taken a first step towards defining ‘experimental secondary ischemia’, a surrogate marker for both early brain injury and delayed cerebral ischemia [80]. Although this definition can help in future primary animal studies, it cannot be retrospectively implemented in previously published studies.

Second striking observation is that delayed cerebral ischemia is measured up to two weeks (336 hours) after ictus as determined by neurological examination. For the late time points (192 to 336 hours) after ictus, it is difficult to say to assess if the models actually display delayed cerebral ischemia. In most of the studies, neurological composite scores indicate that animals are functionally affected, however it remains unclear whether the deficit is caused by the induced subarachnoid hemorrhage or a secondary process in the vein of delayed cerebral ischemia.

Our systematic review shows that there is a significant difference in the terms and definitions, which are used to describe the phenomenon of delayed cerebral ischemia in experimental mouse models. This inconsistency in outcome parameters makes it difficult to compare results between studies, in the same way inconsistency in outcome parameters in human studies has for long obstructed adequate comparison of results. In this study, we assigned all pathophysiological changes and neurological deterioration after six hours following subarachnoid hemorrhage induction to delayed cerebral ischemia. However, this categorization remains disputable.

The discrepancy between experimental and clinical subarachnoid hemorrhage could be another reason for unsuccessful translational results [28]. Experimental subarachnoid hemorrhage and animal models are used to resemble the human condition and to study pathophysiological changes following subarachnoid hemorrhage as well as the effect of pharmaceuticals and interventions. However, these experimental findings might not necessarily mimic the human condition [29]. In humans, delayed cerebral ischemia is exclusively defined by the occurrence of cerebral infarction or clinical neurological deterioration [83]. Moreover, the event rate of delayed cerebral ischemia in most studies was extremely high and nears a 100% of mice. In contrast, the prevalence in humans is around 29% [65]. This adds to the discussion on the scientific value of certain animal models of human diseases as pathophysiological changes after experimental subarachnoid hemorrhage will not completely match human conditions.

Another striking observation was an issue we encountered in the study of Dong et al. [13]. We observed two overlapping histological fields in Figure 4b, in two different panels belonging to two different study groups: *sham* and *sham + vehicle*. We have raised our concerns regarding this issue on Pubpeer ([Link](https://pubpeer.com/publications/47070D29629C93658D707D3B4E1727#1)) and have notified the editor in chief of the *Journal of Pineal Research* in an email. We have not excluded this study from our systematic review as we currently cannot oversee the consequences of these overlapping panels and are awaiting further response from the journal.

There is a correlation between poor reporting in animal studies and poor experimental design, which can introduce bias and exaggerated effect sizes [43,84]. Incomplete reporting was represented in our risk of bias assessment: many items were at an unclear risk of bias because the necessary methodological details to assess the true risk of bias were not reported. This could be improved if authors of primary animals studies followed the Animals in Research: Reporting In Vivo Experiments (ARRIVE) guidelines [61] and if journals and peers enforced the use of these guidelines to improve the reporting quality of animal studies [32]. The large fraction of studies at an unclear risk for most types of bias, makes it very difficult to interpret the results of the included studies.

Finally, we would like to recommend the following to scientists working on experimental mouse models of subarachnoid hemorrhage and delayed cerebral ischemia:

- Register your research protocol, before starting your experiment, in a publicly available database, such as preclinicaltrials.eu. Clearly define aspects of importance such as study population, time points and outcome measures in your protocol. Also define how you will distinguish between injury sustained through the initial subarachnoid hemorrhage and injury sustained through mechanisms such as delayed cerebral ischemia.
- Use ARRIVE writing guideline when preparing your manuscript for publication.
- General recommendations such as using a power calculation to gauge the number of animals needed to demonstrate an effect, or the use of randomization and blinding throughout the study to avoid bias are always important and remain staples of ethical research practices.

## Conclusion

In summary, the results show that established perforation and injection-models for experimental subarachnoid hemorrhage are highly standardized. However, there is a high inconsistency in definitions, which are used to describe the phenomenon of delayed cerebral ischemia in experimental mouse models. Although the perforation model results in a higher rate of delayed cerebral ischemia, there is no effect on the time of occurrence after ictus. It is unclear to what extent the delayed cerebral ischemia demonstrated in these animal models is comparable to the clinical situation.

## Supporting information

Supplement 1 - Search Strategy

Supplement 2 - Data Extraction

Supplement 3 - Risk Of Bias Assessment

## Funding

This study was funded by ZonMw. Grant number: 114024130

## Conflict of interest

Cihat Karadag: none

Igor Fischer, none

Kimberley E. Wever is a PROSPERO administrator (no payment). The PROSPERO record of this review was handled by an independent administrator who is not involved in this review.

On Ying Chan: none

René Aquarius is a PROSPERO administrator (no payment). The PROSPERO record of this review was handled by an independent administrator who is not involved in this review.

Ronald H. M. A. Bartels: none

Jan F. Cornelius: none

Jasper H. van Lieshout: none

Hieronymus D. Boogaarts is a consultant for Stryker neurovascular. Fees are paid to the Neurosurgery department of the Radboud University Medical Center.

Marcel A. Kamp: none

All authors certify that they have no affiliations with or involvement in any (other) organization or entity with any financial interest (such as honoraria; educational grants; participation in speakers’ bureaus; membership, employment, consultancies, stock ownership, or other equity interest; and expert testimony or patent-licensing arrangements), or non-financial interest (such as personal or professional relationships, affiliations, knowledge or beliefs) in the subject matter or materials discussed in this manuscript.

## Availability of data and material

Data available in Supplement 1-3.

